# The diversity of cellular systems involved in carbonate precipitation by *Escherichia coli*

**DOI:** 10.1101/2025.02.06.636833

**Authors:** Matthew E. Jennings, George J. Breley, Reilly S. Blackwell, Anna Drabik, Kathleen R. Gisser, Joe Kainrad, Victoria Ligman, Jaycie Proctor, Rose Deshler, Brian A. O’Hart, Arlyn Rivera, Lorelei Centrella, Sara Bonn, Bisma Chaudhry, Hazel A. Barton

## Abstract

Climate change is increasing the need to limit levels of anthropogenic CO_2_ released into the atmosphere. One approach being investigated is to generate products based on microbially induced carbonate precipitation (MICP), which can trap CO_2_ as CaCO_3_. We recently identified a novel MICP pathway in bacteria that is initiated by Ca^2+^ toxicity in cells, causing extracellular CO_2_ to be trapped as CO_3_ ^2-^ by *Escherichia coli,* although the yield of precipitated CaCO_3_ remained low (in the milligram range). In this work, we used the *E. coli* Keio gene knock-out library to identify 54 genes involved in MICP in *E. coli,* which could be broadly characterized into four groups: central metabolism, iron metabolism, cell architecture, and transport. The role of central metabolism appears to be crucial in maintaining alkaline conditions surrounding the cell that promote CaCO_3_ precipitation. The role of iron metabolism was less clear, although the results suggest that growth rate influences the initiation of MICP. While the impact of repeating polymeric structures on cell surfaces promoting MICP is well established, our results suggest that other structural features may play a role, including fimbriae and flagella. Finally, the results confirmed that Ca^2+^ transport is central to MICP under calcium stress. The results further suggest that the ZntB efflux pump may play a previously unidentified role in Ca^2+^ transport in *E. coli.* By overexpressing some of these genes, our work suggests that there are several previously unidentified cellular mechanisms that could serve as a target for enhanced MICP in *E. coli.* By incorporating these processes into MICP pathways in *E. coli,* it may be possible to increase the volume of CO_2_ fixed using this pathway and yield potentially new products that can replace CO_2_ intensive products, such as precipitated calcium carbonates (PCCs) for industry.

## Introduction

Anthropogenic emissions of greenhouse gases leading to increasing global temperatures represent one of the greatest threats to our biome, with impacts including an increase in extreme weather events, reduced biodiversity, and damage to important ecosystem services [1–5]. The push to find solutions to remove excess anthropogenic CO_2_ include a range of geological and biological sequestration approaches, such as deep burial and forestry management [6, 7]. There is also an increased interest in utilizing carbonatogenesis by microorganisms, which allows the capture of carbon dioxide as CO_3_ ^2-^, which is subsequently precipitated as CaCO_3_ to prevent release back into the atmosphere [8, 9]. Such microbially driven carbonate precipitation has been adapted to various biotechnology applications, including cement-based materials, algal biofuels, and soil amendments [10–14].

Biologically induced mineralization (BIM) occurs when an organism indirectly causes the precipitation of minerals by altering the chemistry of the surrounding environment through metabolic activity, or acts as physical nucleation points for crystal formation [10]. Carbonatogenesis resulting from these processes is known as microbially induced CaCO_3_ precipitation (MICP), which can occur through a variety of pathways, including ureolysis, photosynthesis, sulfate reduction, nitrate reduction, ammonification, and methane oxidation [15]. Metabolic activity in MICP often alters the local pH, which shifts the dynamic equilibrium of bicarbonate in solution to favor CO_3_ ^2-^ production. This causes them to exceed their saturation index (SI) and precipitate with a divalent cation (usually environmentally abundant Ca^2+^) [16]. Most MICP used in industrial applications use microbial ureolysis, where urease cleaves urea into CO_2_ and 2NH_4_^+^; as a weak base, NH_4_^+^ increases the extracellular pH, driving the precipitation of CaCO_3_ [17]. This urease-dependent method has been favored in industry due to its ease of use and cost effectiveness, although limitations include the availability of urea and an unpleasant odor [18, 19].

Particulate (μm scale) calcium carbonate (PCC) has a number of industrial uses: in paper, carbonates are used as a filler and coating, which can increase the smoothness, brightness and help preserve paper; in thermoplastics, as a filler to reduce polymer volume and modulate elasticity; in sealants and adhesives as a thixotropic agent to reduce shrinkage as polymers set; and in coatings, where carbonates play an important role in opacity, brightness, gloss and durability [20–22]. Over 130,000 kilotons of PCCs were produced worldwide in 2019 in two primary forms [23]: 1) ground calcium carbonates from the mining of calcitic rocks (limestone, marble, travertine, and chalk); and 2) chemically precipitated PCC from CaO (produced by calcination at >1,000°C) reacted with CO_2_. Both approaches require the mining of calcitic rock, with an increasing pressure on ecosystems and the groundwater found within these landscapes, while the production of PCCs contributes to significant global CO_2_ emissions [24, 25].

Recently, we were able to demonstrate PCC production in liquid cultures of *Escherichia coli* that sources the CO_3_ ^2-^ from atmospheric CO_2_, via a novel calcium stress-dependent MICP mechanism first described in microorganisms from calcitic cave environments [26]. This pathway relies on the use of potentially toxic levels of Ca^2+^ via growth on media with a calcium carboxylate salt (calcium acetate, calcium propionate, etc.) [27]. These PCCs have the potential to sequester significant amounts of atmospheric CO_2_, but also remove the need for energy intensive methods in their production and transport, and would theoretically sequester >1-ton CO_2_/ton metabolic PCCs (mPPCs) produced [25].

While there are numerous industrial applications for a mPCC product, the current yield of PCC from *E. coli* cultures is low (∼100 mg PCC/ 25 mL) containing 62.5 g of Ca^2+^ ions. We therefore decided to investigate whether CaCO_3_ precipitation in *E. coli* could be enhanced by genetic modification using the Keio collection; this strain library contains non-polar mutations in 4,000 non-essential *E. coli* genes [28]. By screening the Keio library for genes that affected precipitation under calcium stress on calcium propionate, we identified over 50 potential genes involved in a myriad of cellular activities, including central metabolism, cell structure, and transport. The data confirmed that the ability to utilize the organic calcium salt is critical to calcium-stress MICP, while central metabolism plays an important role in carbonatogenesis, from limiting the acidic products of fermentation and cellular CO_2_ levels. While PCC production in *E. coli* increases the range of industrial applications where products for carbon sequestration could be used, it remains unclear whether any of the genetic modifications identified could produce the high levels of CaCO_3_ necessary to scale these green PCCs as an industrial replacement to those currently in use.

## Materials and Methods

### Bacterial strains and growth conditions

*E. coli* strain BW25113 [29] or K-12 were used as the wild type strain when assaying the various carbonate precipitation phenotypes. The strains were maintained on LB agar plates at 37°C. Single gene deletion strains were obtained from the Keio collection, and consist of single gene mutations in the BW25113 background strain [28]. Single gene deletion strains were maintained on LB agar plates with 50 μg/mL kanamycin at 37°C.

The precipitation of CaCO_3_ was assayed on modified B4 media. Standard B4 media contains 4 g yeast extract, 10 g glucose, and 15 g agar per liter of media adjusted to pH 7.2 before autoclaving [30]. We used a minimal B-4 formulation (B-4m), which was prepared as B4 media, but glucose was omitted, with 2.5 g of the relevant calcium salt dissolved in water and filter sterilized before addition to the media following autoclaving. The calcium sources used were calcium acetate (B-4m), calcium propionate (B-4mPr), calcium L-lactate hydrate (B-4mLa), calcium succinate monohydrate (B-4mSu), and calcium pyruvate (B-4mPy). To improve the solubility of calcium succinate, it was dissolved in 0.1M HCl and readjusted to pH 7.2 before filtering. All solid media was prepared with 1.5% agar. To measure the change in pH in the media upon growth, a single line of WT, *ΔnuoA*, *ΔsdhC*, *ΔatpF*, and *ΔprpD* were streaked onto separate media plates and incubated at room temperature for 24 hours. The pH was measured using an Orion pH meter with an internal reference probe (Fisher 9863BN) inserted in the agar directly adjacent to the line of growth in three separate locations.

To assay the Keio collection for differences in carbonate precipitation, strains were plated onto B-4mPr agar. Media was poured into sterile Nunc OmniTray dishes (Thermo Scientific, Rochester, NY) and allowed to solidify. A 96-pin inoculator was used to spot inoculate 96 strains onto a single plate. Plates were incubated one week at room temperature, and then examined for carbonate precipitation using an Olympus BX53 light microscope (Olympus Life Science, Center Valley, PA) at 100x magnification. Carbonate precipitation was qualitatively compared to WT (BW25113) to determine if strains displayed enhanced or deficient carbonate precipitation phenotypes.

The iron amendment experiments were established in 100 mL B-4mSu in a 250mL Erlenmeyer flask. The media was inoculated with 100 μL of an overnight culture of *Escherichia coli* MG1655 (K-12; American Type Culture Collection #700926) and stirred continuously on a Corning PC410-D stir plate with a 3 cm stir bar at 170 rpm. The pH was logged continuously using a REED R3000SD pH meter with a REED R3000SD-PH2 General Purpose pH Electrode. Prior to each experiment, the electrode was calibrated according to the manufacturer’s instructions, sterilized by immersion for twenty minutes in 1M HCl, and then washed in sterile DI water.

### Cloning and gene expression

Genes were amplified from *E. coli* strain K-12 using standard techniques. Primers were designed using the design primer tool in Geneious Prime 2022.0 (GraphPad Software, Boston, MA), to avoid self-annealing sequences. Forward primers were designed to contain an EcoRI site at the 5’ end, and reverse primers contained an XbaI site at the 5’ end to facilitate cloning. All primers used in this study were purchased from IDT. A list of primers used for amplification of the native *E. coli* genes can be found in S1 table PCR was carried out using Phusion high fidelity polymerase (New England Biolabs, Ipswich, MA) and the PCR products were purified using a Wizard SV PCR Clean-up Kit (Promega, Madison, WI) and quantified using a NanoDrop spectrophotometer (Thermo Fisher, Waltham, MA). Each PCR fragment was cloned into plasmid pPRO24 [31], which contains a propionate driven promoter (P_prpB_). Plasmid and PCR products were both digested with EcoRI and XbaI restriction enzymes (New England Biolabs) prior to ligation, and purified as described above. Ligation products were used to transform *E. coli* strain DH5α (Thermo Fisher), and successful transformants were sequenced to confirm the presence of each gene without mutations. Sequencing primers are listed in Table S1. Plasmids were then used to transform *E. coli* DE3 cells (Sigma Aldrich, St. Louis, MO) to generate expression strains. Strains containing the plasmid were maintained in the presence of 100 µg/mL ampicillin. Induction of gene expression on plates was achieved by adding 2.5 g/L Na-propionate to the media. Induction of gene expression in liquid media was achieved by adding 50 mM Na-propionate, as not to confound the CaCO_3_ precipitation experiments.

### Quantification of carbonate precipitation in liquid media

Quantification of solid CaCO_3_ precipitates in liquid media, 5 mL cultures of carbonate precipitation media were inoculated with 60 µL of an overnight culture of *E. coli* strains grown in LB. The cultures were incubated 24 hours at 37℃. The cultures were harvested by spinning at 5,183 x *g* in an Avanti J-E centrifuge for 10 minutes. The supernatant was removed, and pellets washed 1X in 1 mL 1x PBS pH 8. The pellet was resuspended in 1 mL of 5% HNO_3_ to dissolve any precipitated calcium carbonate for 30 minutes at room temperature [26]. The supernatants were assayed using the Arsenazo III calcium sensitive dye (Pointe) at 650 nm on a Molecular Devices Spectramax M4 spectrophotometer. A standard curve was generated using soluble calcium (Inorganic Ventures 10PPM) at 4, 6, 8, 10, 12, and 15 mg/dL. There was no significant difference in CaCO_3_ production between *E. coli* BW25115 and DE3 in this assay.

## Results

### Identification of the genes involved in carbonate precipitation

To identify genes associated with CaCO_3_ precipitation, we spot-plated knockout strains from the Keio *E. coli* library [28] onto a calcium propionate medium lacking glucose (B-4mPr) and monitored for 1 week at room temperature. We previously demonstrated that this media is an effective screen for carbonatogenesis in *E. coli* [26], with the formation of visible CaCO_3_ crystal after 4 days at room temperature [26]. To demonstrate the effectiveness of this approach, the screen identified *prp*D, which functions in the 2-methylcitrate pathway of propionate catabolism, and cells lacking this gene cannot utilize calcium propionate for growth (Fig 1) [32]. Other genes involved in propionate catabolism (*prpB* and *acnB*) were identified to have a modest effect on precipitation, compared to *ΔprpD* (data not shown). Of the ∼4,000 strains screened, we observed 54 mutants with a WT growth phenotype, but altered CaCO_3_ precipitation phenotype. This included a variety of known cellular functions that could be clustered into 4 major groups: central metabolism, iron metabolism, cell architecture, and transport (Table 1). Eleven additional genes with miscellaneous cellular functions were also identified, including the *prp*D control. One cytoplasmic membrane protein of unknown function (*ygd*Q) was identified, which was not investigated further (Table 1). All knockout strains that were identified were re-streaked to confirm the loss of the carbonatogenesis phenotype.

**Fig 1.**
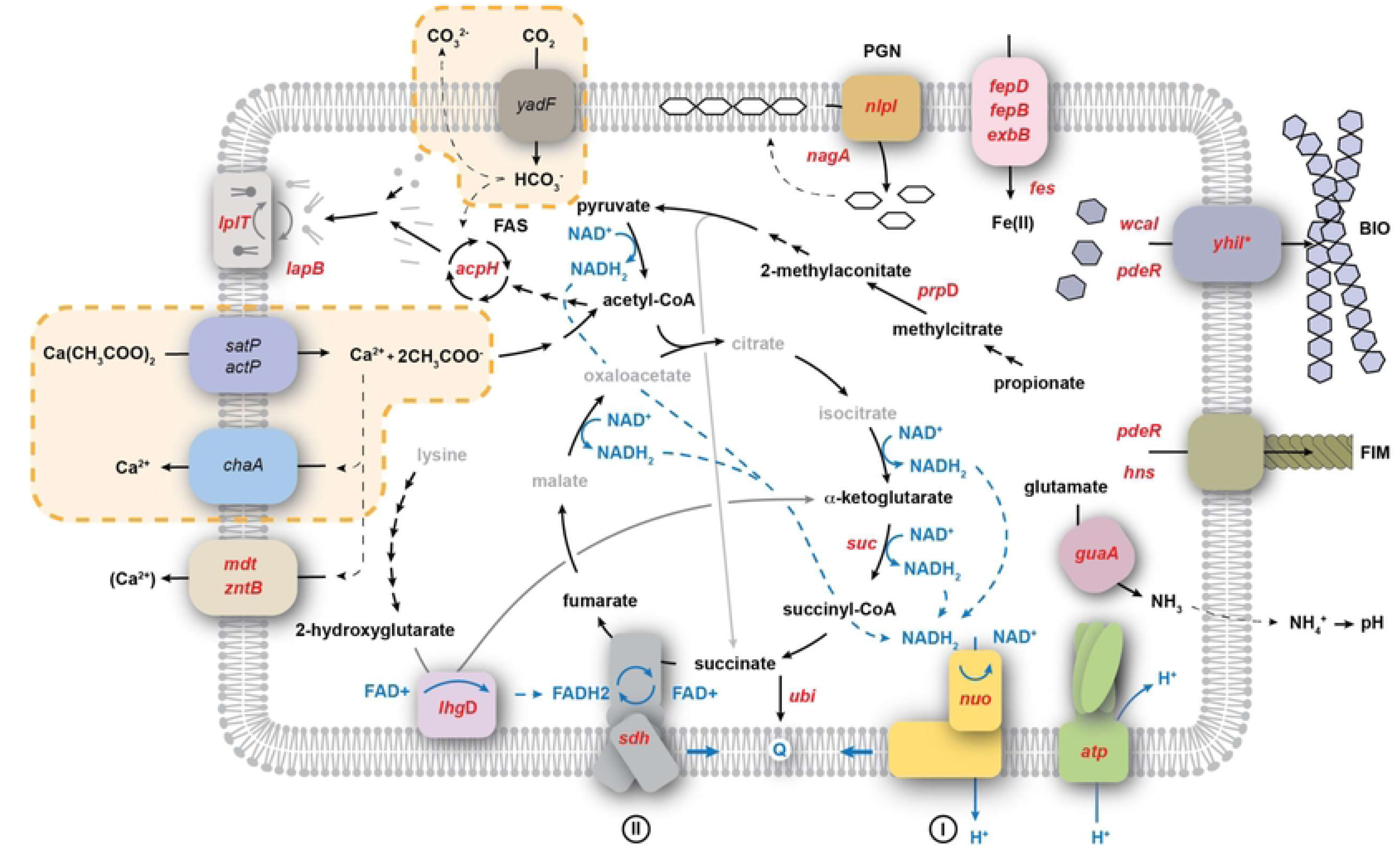
Metabolic pathways influencing carbonatogenesis identified in this study. The calcium homeostasis pathway that drives carbonatogenesis in our model is highlighted in the yellow dashed boxes: uptake of the organic calcium salt (represented by calcium acetate) via the sat or act operons leads to the metabolism of the carboxylic acid (the glyoxylic acid pathway for acetate metabolism within the TCA is not shown for simplicity), with the excess Ca^2+^ removed via an unknown calcium transporter. The uptake of CO_2_ via the carbonic anhydrase (*yadF*) provides the source of CO_3_ ^2-^ ions for precipitation. The Keio library knockout strains identified with defects in carbonatogenesis are shown in red. While ChaA was previously demonstrated to be the Ca^2+^ transporter in *Salmonella* [27], our screen indicated a phenotype for the transporters *mdtH*, *mdtL*, and *zntB*. The intermediates shown in central metabolism and metabolism of the provided calcium sources are shown in black (central metabolites not identified with a role in carbonatogenesis in this work are gray). Blue indicates the flow of electrons and reducing equivalents. The cell wall and membrane structure of *E. coli* is represented as a single membrane for simplicity.

**Table 1.**
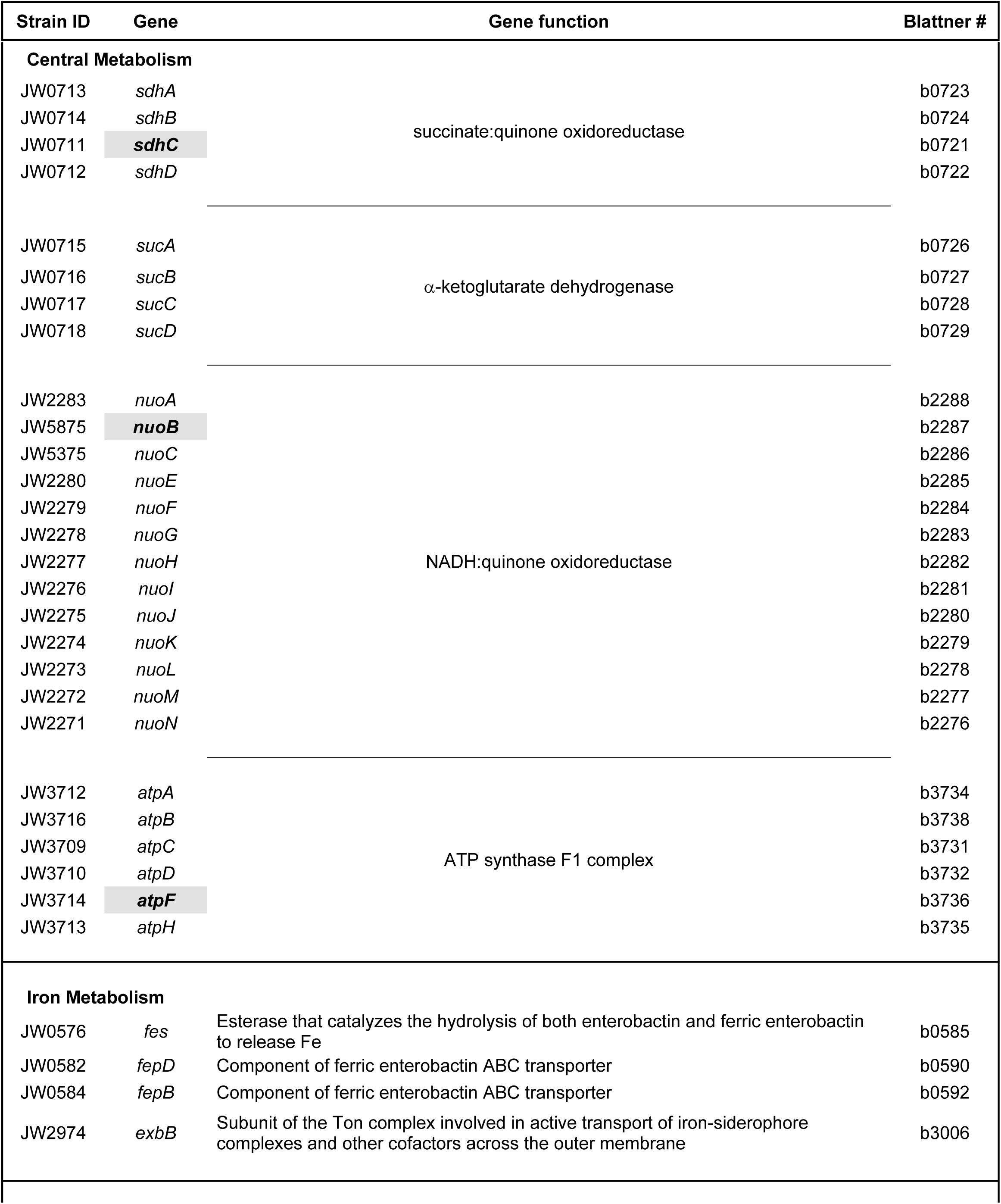
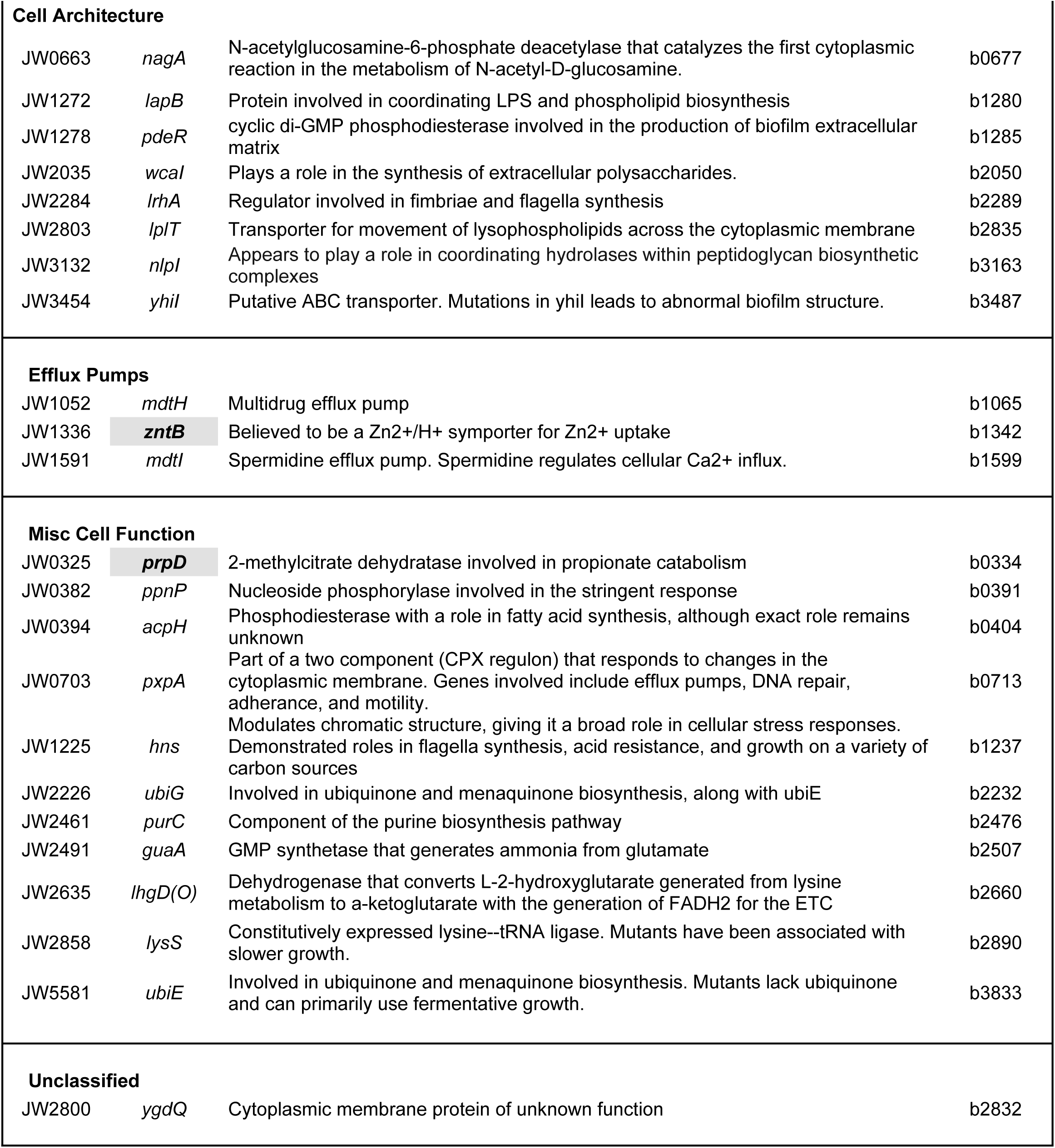
List of *E. coli* Keio knockout strains with altered CaCO_3_ precipitation phenotype.

### The role of central metabolism

Of the genes identified in central metabolism, all were associated with protein complexes in the tricarboxylic acid (TCA) cycle and electron transport (Table 1 and Fig 1): succinate:quinone oxidoreductase, SQR (*sdh*CDAB); α-ketoglutarate dehydrogenase (*suc*ABCD); NADH:quinone oxidoreductase, Complex I [*nuo*ABC(D)EFGHIJKLMN]; and ATP synthase (*atp*ABCDFH). The fact that every representative gene from each of these operons was identified in our screen of the Keio library confirms the role that these pathways play in carbonatogenesis on propionate (Table 1). The lack of major components of the TCA cycle and electron transport can affect metabolism in a variety of ways, so we decided to examine whether the observed effects were related to growth rate or specific to carbonatogenesis, and narrowed down our experiments to the representative genes, Δ*nuo*B, Δ*sdh*C, and Δ*atp*F, using WT and Δ*prp*D as controls (Table 1). In the case of α-ketoglutarate dehydrogenase, mutations in the *suc* operon have been shown to reduce the amount of CO_2_ released by the TCA cycle by >50% [33]. Given the need to create CO_3_ ^2-^ ions for CaCO_3_ precipitation, we felt that it was not necessary to explore the impact of the *suc* knockouts further.

The use of solid media allows us to rapidly assess CaCO_3_ precipitation by visual inspection, but it also allows us to examine some of the local conditions that are difficult to measure in liquid culture, such as the distribution of pH gradients within the environment [26]. We have previously shown that growth of WT *E. coli* in B4-m media produces basic conditions, regardless of the calcium source, presumably through the consumption of amino acids and export of NH_3_ into the surrounding media [26]. When the pH approaches 8.3, the ion activity product exceeds the SI for Ca^2+^ and CO_3_ ^2-^, leading to CaCO_3_ precipitation [26]. As has been previously shown, on all of the calcium sources used, WT *E. coli* raised the pH (range 8.1 – 8.5) and led to the precipitation of CaCO_3_ (Fig 2A) [26]. The cells need to catabolize the organic calcium salt for carbonatogenesis (Fig 1) [27], and unsurprisingly, the Δ*prpD* knockout only prevented CaCO_3_ precipitation on calcium propionate (Fig 2A).

**Fig 2.**
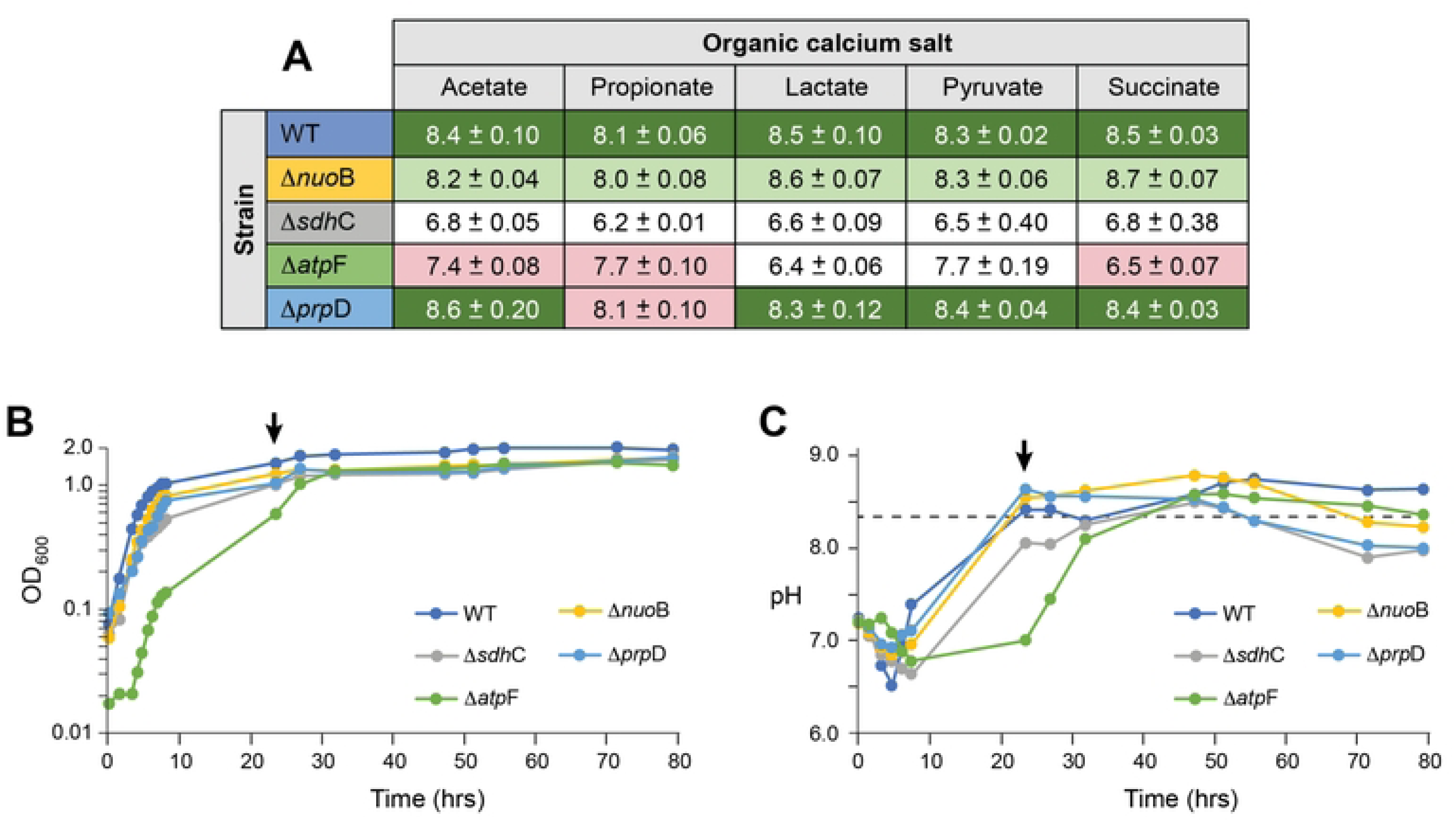
Qualitative comparison of calcium source on carbonatogenesis in selected *E. coli* knock-out mutants compared to WT. A) Strains were spot plated onto carbonatogenesis media containing the indicated organic calcium-salt. The pH of the media after 24 hours of growth at 37°C was measured (the average of three pH measurements +/− SD). The color of the cell is based on a visual qualification of the amount of PCC produced relative to WT; deep green, high precipitation; pale green, limited precipitation, white, no precipitation; pink indicates the knock-out prevents use of the indicated organic calcium salt for growth. B) The growth rate in liquid culture as measured by OD_600_. The initiation of carbonate precipitation can be observed by a bump in OD_600_ (indicated in WT by the black arrow). C) Change in the pH of the media corresponding to the liquid culture in (B). A drop in pH indicates the initiation of carbonatogenesis (indicated in WT by the black arrow).

The loss of *sdhC* prevents the assembly of the SQR complex, which catalyzes the oxidation of succinate to fumarate coupled to the reduction of FAD in the TCA cycle [34]. The lack of this enzyme prevents proper functioning of the TCA cycle; however, the enzyme fumarate dehydrogenase, which catalyzes the reduction of fumarate to succinate, can operate in the reverse direction, albeit at reduced efficiency with secretion of acetic acid [35, 36]. Our data indicated that the *ΔsdhC* knockout acidifies the surrounding medium under all conditions tested (Fig 2A), presumably through the production of this acid. Growth in liquid cultures allows us to qualitatively evaluate the growth kinetics of each strain, and when carbonatogenesis occurs it can be observed by a notable increase in the OD_600_ signal (via light scattering by the formation of CaCO_3_ particulates) and a simultaneous drop in pH (as CO_3_ ^2-^ is consumed) [26]. These data (Fig 2B) indicate that the overall growth rate of *ΔsdhC* was slower that WT (mid-log doubling time 1.7 hrs, compared with 1.27 hrs for WT), with the pH only minimally exceeding the SI for CaCO_3_ after 30 hrs of growth (Fig 2C).

The AtpF protein is involved in the formation of the ATP synthase complex, requiring growth on fermentable carbon sources [37, 38], and the Δ*atpF* strain did not produce carbonate regardless of calcium source (Fig 2A). Lactate and pyruvate can be fermented, which would presumably lead to the production of acetic and other acids [39–41]; however, there is no clear difference in the pH of the surrounding media between the use of fermentable (lactate and pyruvate) and non-fermentable calcium sources (acetate, propionate, and succinate) in the media (Fig 2A). The presence of a lag phase and much slower growth for Δ*atpF* likely corresponds to the shift in cellular metabolism from respiratory to fermentative growth (Fig 2B), and given that the Δ*atpF* strain produced colonies on non-fermentable calcium acetate, propionate, and succinate, it suggests that the observed growth is dependent on the amino acids present in the media (as for the Δ*prpD* knockout on propionate; Fig 2A). Nonetheless, while growth in the Δ*prpD* led to an increased pH in the surrounding media, Δ*atpF* did not, suggesting a more complex response to the formation of basic conditions during amino acid catabolism (Fig 2A and C).

### The role of central metabolism – Complex I

The strain Δ*nuoB* had reduced levels of CaCO_3_ precipitation in our initial screen (Table 1). While this mutant raised the pH using all calcium sources tested (pH range 8.0 – 8.7), it maintained a reduced precipitation phenotype on all media (Fig 2A). Given the critical role of Complex I in electron transport, it was possible that this knockout was simply affecting the growth rate, which in turn would reduce catabolism of the calcium salt and observed levels of CaCO_3_ precipitation [26, 27], but while Δ*nuoB* grew more slowly in liquid culture, it was more comparable to WT than other assayed mutants (mid-log doubling time 1.51 hrs; Fig 2B).

We decided to explore the slow onset of carbonatogenesis in Δ*nuoB* in a more quantitative way and assayed the amount of CaCO_3_ precipitated between Δ*nuoB,* WT, and Δ*prpD* in liquid cultures containing the various calcium sources (Fig 3). The amount of CaCO_3_ in the cultures was measured at 24 hr, when the initiation of precipitation is observed in WT (Fig 2C). The results demonstrated limited precipitation in acetate and propionate between Δ*nuoB,* WT, and Δ*prpD* at 24 hrs, which is consistent with prior observations of a lower level of carbonatogenesis in *E. coli* using these calcium sources [26]. When grown on calcium lactate, pyruvate, and succinate, Δ*prpD* demonstrates the same precipitation phenotype as WT, with the highest amount CaCO_3_ precipitation when these strains were grown on calcium succinate (Fig 3). The data also demonstrated that Δ*nuoB* precipitated significantly less CaCO_3_ than either WT or Δ*prpD* on calcium lactate, pyruvate, and succinate (Fig 3). These data correlate with the quantitation results seen on the solid media, confirming the effectiveness of the spot plating method as a rapid screen to observe genetic impacts on carbonatogenesis in *E. coli* (Fig 3).

**Fig 3.**
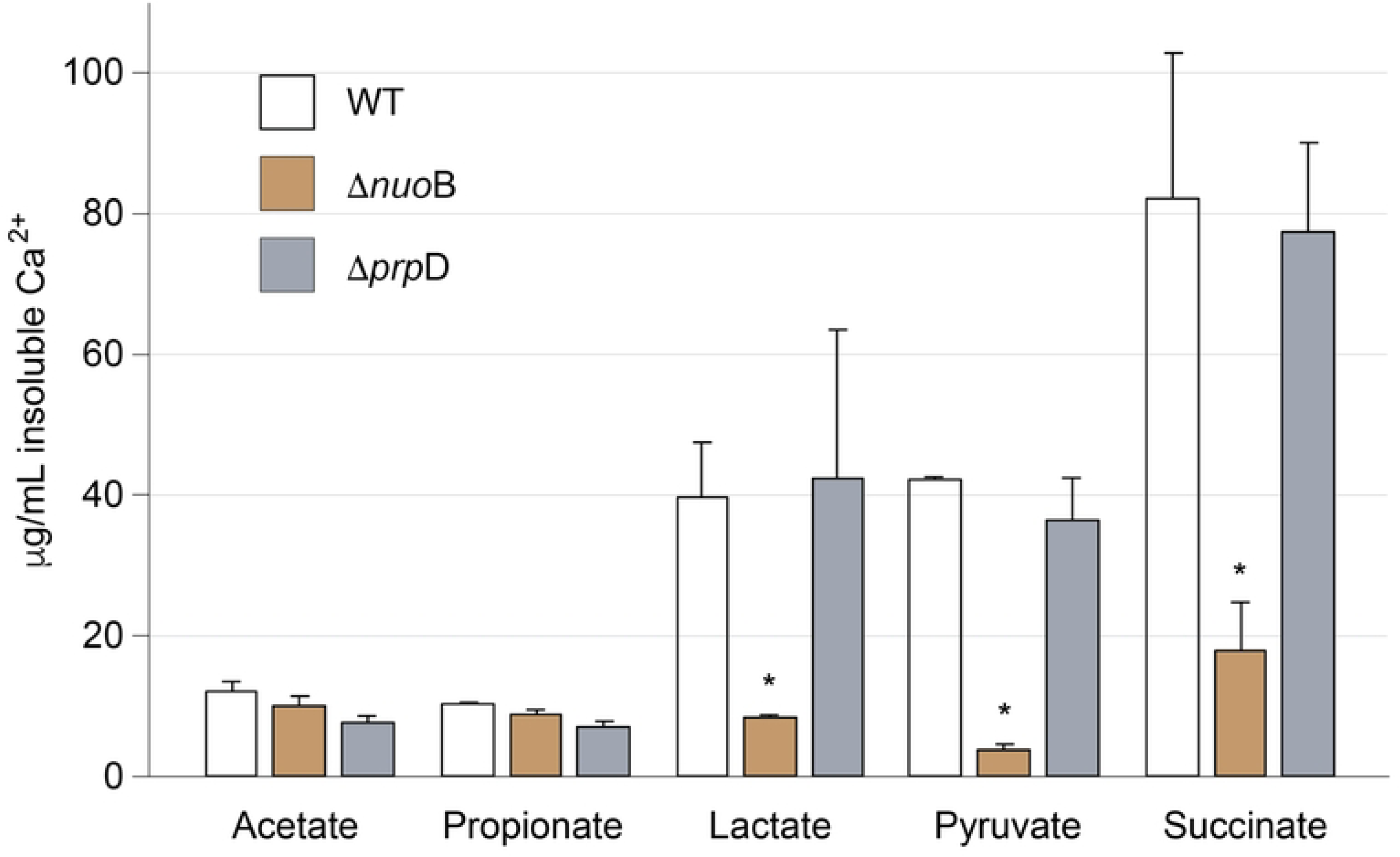
Quantification of insoluble Ca^2+^ from 25 mL B-4m media containing the indicated calcium salt following 24 hr incubation. Ten mL of each culture was harvested, soluble Ca removed via washing, and then the remaining pellet treated with 5% HNO_3_ to dissolve CaCO_3_. Values are average of three replicates and error bars represent standard deviation. An asterisk represents a significant difference compared to the BW25113 values (WT) using a Student’s T-test (p < 0.05).

### Iron metabolism

Several genes involved in iron acquisition and transport were identified as affecting carbonatogenesis (Table 1). The relationship of iron acquisition and transport genes to CaCO_3_ precipitation is not clear, though the issue may simply be poor growth due to limiting iron when these genes are missing [42]. To determine if the precipitation phenotype is related to iron transport, we examined the impact of amending cultures with iron. We continuously monitored pH as a proxy for the initiation of CaCO_3_ precipitation (Fig 4), with a sudden drop in pH corresponding with the onset of CaCO_3_ crystal accumulation in the bottom of the flask (data not shown) [26]. We performed this assay at room temperature, to slow the rate of growth and provide a better resolution of the timing of carbonatogenesis.

**Fig 4.**
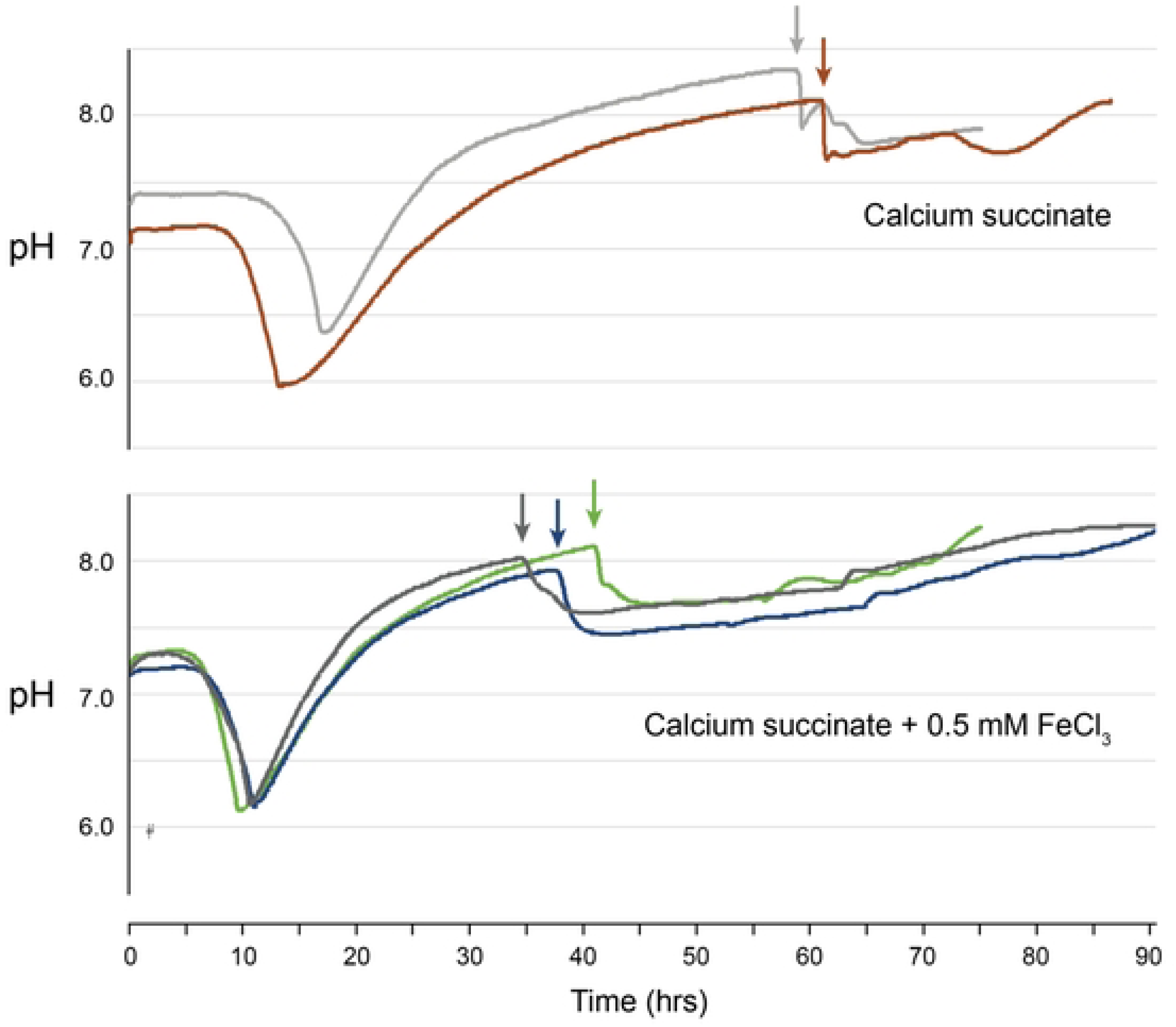
The impact of iron on CaCO_3_ precipitation rate. Cultures were grown at room temperature in calcium succinate, with pH used as a proxy for the initiation of CaCO_3_ precipitation. The pH of the culture was monitored continuously, with and without the addition of 0.5 mM FeCl_3_. Black arrows indicate the initiation of carbonatogenesis.

When WT *E. coli* was grown using calcium succinate, precipitation began around 60 hrs (Fig 4), which corresponds with the observations of similar culture conditions [26]. When the media was amended with 0.5 mM FeCl_3_, the onset of precipitation was reduced to between 32 – 41 hrs, without a significant change in growth rate (Fig 4), suggesting that Fe(III) amendment does influence the timing of carbonatogenesis. The resulting crystals had a distinctive brown color, suggesting that available Fe(III) was being incorporated into the growing crystals (data not shown).

### Cell architecture

Our screen similarly suggested that a variety of extracellular structures in *E. coli* influence CaCO_3_ precipitation (Table 1). Bacteria bind metal ions, such as Ca^2+^ by virtue of the acid/base properties of the cell wall functional groups, which include amino, carboxylic, hydroxyl, and phosphate sites [43, 44]. The genes identified in the Keio library screen suggests that there is a myriad of factors that may influence this coordination and nucleation of the forming CaCO_3_ nanocrystals, including EPS, LPS, and peptidoglycan (Table 1), which have previously been demonstrated to play an important role in nucleating CaCO_3_ crystals. Nonetheless, our data also identified the potential for the proteinaceous repeating units of fimbriae and flagella in promoting such nucleation, although the experiments needed to study these interactions are beyond the scope of this study.

### Efflux pumps

Our screen identified three other transporters, *ΔmdtH, ΔmdtI,* and *ΔzntB* (Table 1), which have not previously been associated with Ca^2+^ transport [45]. The *mdtH* (*yceL*) and *mdtL* (*yidY*) transporters were identified through sequence annotation of the *E. coli* genome as members of the major transporter superfamily involved in antibiotic resistance, although the over expression of *mdtH* or *mdtL* only weakly enhanced resistance [45, 46]. No additional work has attempted to understand the cellular function of these transporters [45]. The ZntB transporter protein is believed to be involved in zinc homeostasis, although whether it functions as an importer or exporter remains under debate [47, 48].

Given the greater level of understanding of *zntB* function (compared to *mdtH* and *mdtL* [47]), we decided to study its effect on carbonatogenesis by cloning into an expression vector (pPRO24) in *E. coli*, which includes a promoter under the control of the propionate-inducible prpR regulator [31]. We also cloned *yrbG,* which encodes a sodium/calcium ion exchanger that regulates *E. coli* cytosolic calcium, but has no described role in MICP [49]. Along with *zntB,* both genes were inserted into pPRO24 and sequenced to confirm that they had 100% identity to the WT gene (data not shown).

The expression plasmids (pPRO::*yrbG* and pPRO::*zntB*) were transformed into *E. coli* DE3 and induction of the target proteins, YrbG and ZntB, was confirmed by SDS-polyacrylamide gel electrophoresis (SDS-PAGE; data not shown). We then assayed these constructs, along with a plasmid control, for CaCO_3_ production with or without induction with sodium propionate (Fig 5). There was a small, but not significant, increase in CaCO_3_ precipitation in the plasmid control with the addition of the inducer (25 mM sodium propionate; Fig 5), which may be due to a faster growth rate from the metabolism of supplemented sodium propionate (Fig 5). The precipitation profile of the pPRO::*yrbG* demonstrated a small increase in CaCO_3_ precipitation without induction, but this was not significantly different from WT (with or without induction; Fig 5). We did see a significant impact on the precipitation phenotype in pPRO::*zntB,* with a 3-fold increase in CaCO_3_ precipitation in both the uninduced and induced strain (Fig 5). The impact of induction of *zntB* with sodium propionate was not significant; however, the pPRO24 promoter is known to be leaky, allowing a potential dosing effect from the multicopy plasmid without induction [31].

**Fig 5.**
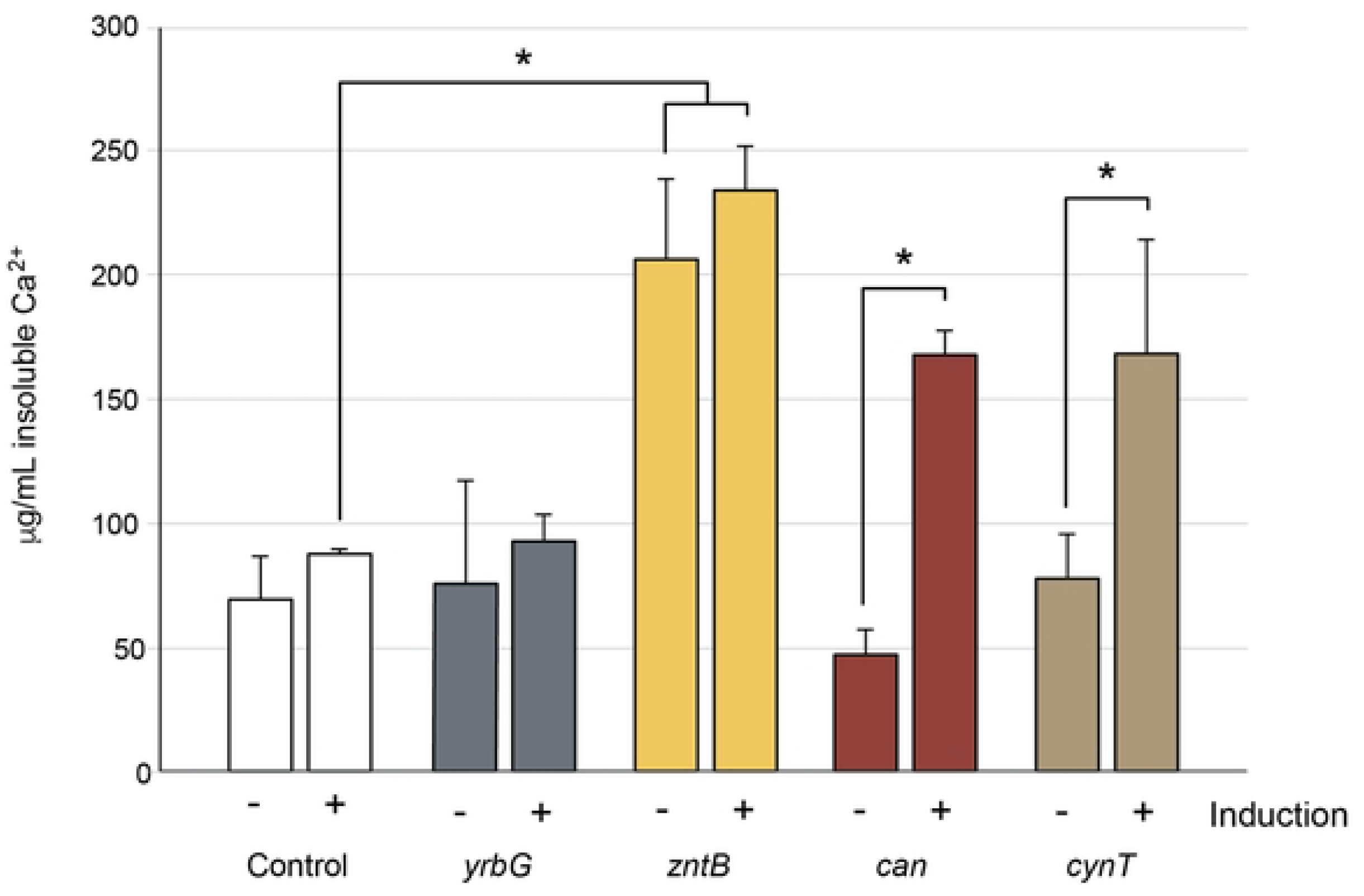
Quantification of CaCO3 precipitation from genetic constructs in *E. coli* DE3. Cells containing the plasmid control (control) or calcium transporter (*yrbG*; which does not play a role in carbonatogenesis) were compared with the transporter *zntB*. Cultures were either uninduced (−) or induced (+) with 25 mM sodium propionate. To confirm that our assay did induce enhanced carbonatogenesis through induction, two genes known to increase CaCO_3_ precipitation (can and cynT) we used. Each bar represents the average of three replicate cultures, with bars corresponding to standard deviation. Statistical comparisons were done using a Student’s T-test, with a significance of p <0.05 indicated by an asterisk.

Prior research has demonstrated an important role for CA in CaCO_3_ precipitation in bacteria, including a periplasmic expressed CA in *E. coli* [50–53]. These genes convert atmospheric CO_2_ to bicarbonate and their overexpression has previously been demonstrated to dramatically improve carbonate precipitation [52]. To compare the impact of CA expression with the *zntB,* we cloned the *E. coli* CA genes *can* (*yadF*) and *cynT.* Upon induction, a significant increase in CaCO_3_ precipitation was detected, which is consistent with previous studies, and demonstrated the validity of our expression approach [52, 54].

### Miscellaneous cell function

A variety of other genes were identified in our knockout assays (Table 1), with a range of cellular functions. While the role of some of the genes in CaCO_3_ precipitation may be relatively straightforward (for example, *guaA* releases ammonia from the catabolism of amino acids and raises pH), others may be more challenging to explain. In particular, there were a number of genes involved in the cellular stress response (*ppaP, pxpA, hns*). It is our understanding that the calcium homeostasis phenotype leading to CaCO_3_ precipitation is driven to some extent by cellular growth under non-ideal conditions, which may require the cell stress response to maintain active growth [27]. Understanding the role of these genes will require additional investigation beyond the scope of this work.

## Discussion

MICP is an exciting developing area of applied microbiology, both as a potential avenue of carbon sequestration and a way to enhance building materials for self-repair [16]. The most common MICP approach in building materials, such as cements, is ureolysis [17]. But the release of ammonia can lead to odor issues, while ammonia can be oxidized to N_2_O, itself is a greenhouse gas [55]. We previously demonstrated carbonate precipitation by *E. coli* using calcium homeostasis (Fig 6) [27], and in this paper we aimed to identify other genes that could enhance this pathway for industrial applications independent of ureolysis. Our data indicated that there are a wide variety of cellular functions that influence MICP in *E. coli*, from central metabolism to cell structure and the stress response (Table 1). While the specific function of many of these genes still needs to be elucidated, they did provide some new clues to the metabolic pathways that influence CaCO_3_ precipitation in *E. coli* and allowed us to refine our model of how cellular Ca^2+^ stress drives carbonatogenesis (Fig 6).

**Fig 6.**
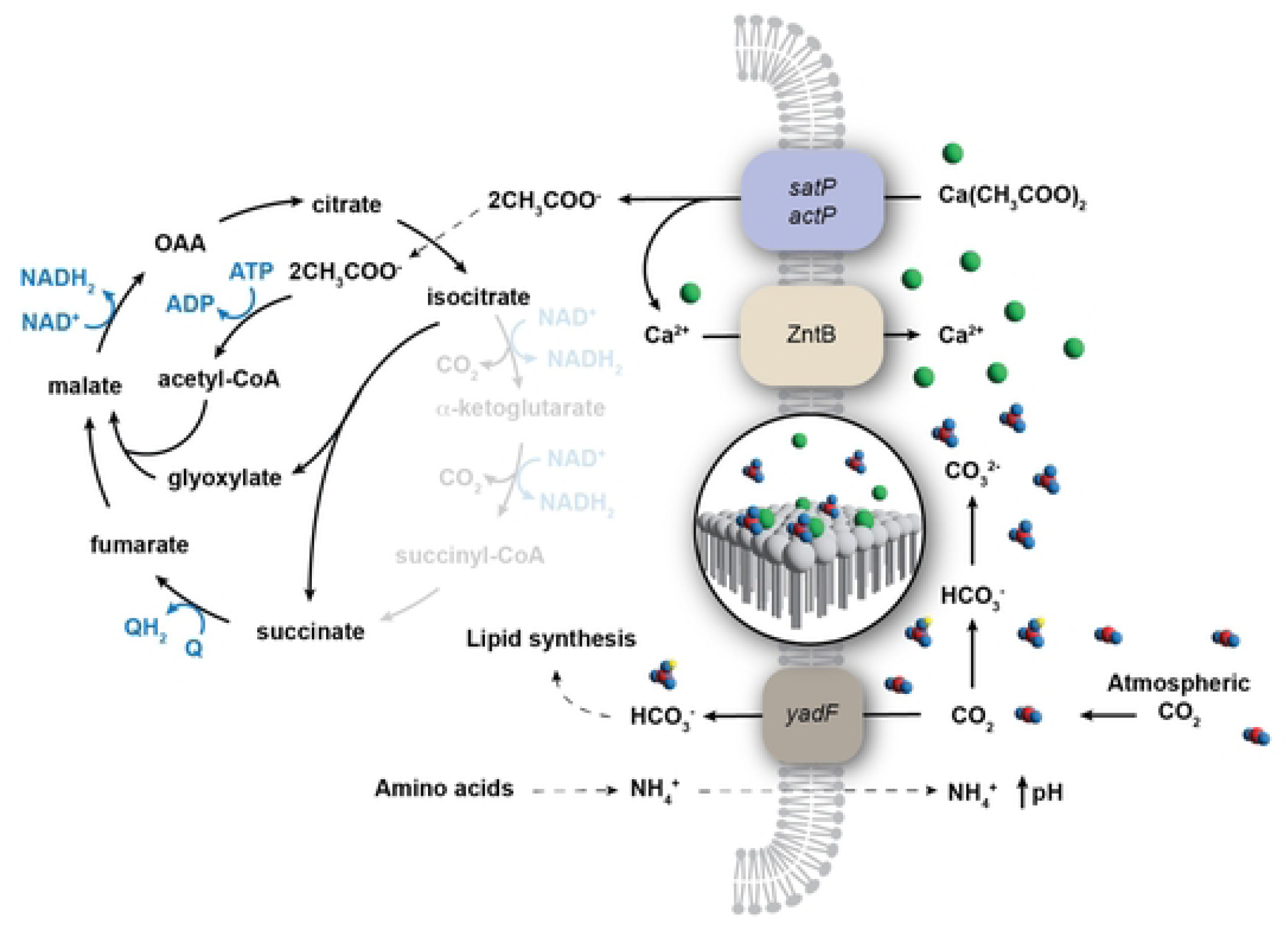
Revised model for carbonatogenesis via calcium homeostasis. The initial steps of our revised carbonatogenesis pathway via calcium homeostasis remain unchanged: an organic calcium salt (represented by calcium acetate) is taken up by the cell via the *sat* or *act* operons which leads to the metabolism of the carboxylic acid via the glyoxylate shunt of the TCA cycle. Carbonic anhydrase (*yadF/can*) produces bicarbonate from atmospheric CO_2_ for biosynthetic purposes to compensate for the loss of CO_2_ from central metabolism (the missing portions of the TCA cycle are indicated with light shading) via the glyoxylate shunt. The update of CO_2_ leads to a reduction in extracellular H_2_CO_3_, raising the surrounding pH. Our data has shown that in *E. coli,* ZntB is the likely exporter of excess Ca^2+^ ions, which bind to negatively charged surface of the cell. The coordination of these Ca^2+^ ions reduces the energy necessary to initiate mineralization. The catabolism of amino acids leads to the release of NH_4_ ^+^, which additionally raises the pH of the surrounding environment, increasing the relative ratio of CO_3_ ^2-^ ions surrounding the cell. Combined these conditions cause the local environment surrounding the cell to favor the precipitation of calcium carbonate, with CO_2_ taken up in both cellular biomass and the growing carbonate minerals. Blue indicates the flow of electrons and reducing equivalents. The cell wall and membrane of *E. coli* is represented as a single membrane for simplicity.

Carbonatogenesis via Ca^2+^ homeostasis requires the organic calcium salt to be taken up by the cell and metabolized (Fig 6) [26, 27]. Our screen on B-4mPr media (Table 1) identified methylcitrate dehydratase (*ΔprpD*), which is involved in the catabolism of propionate, indicating the robustness of our screening approach. But there are four other genes involved in the 2-methylcitrate cycle, which converts propionate into succinate and pyruvate (*ΔprpB, ΔprpC, ΔprpE* and *ΔacnB*), which were not identified in our screen (Table 1) [56]. The genes *ΔprpB* and *ΔacnB* function downstream of *prpD,* and a knockout in all three genes would lead to an increase in cellular levels of 2-methylcitrate, a co-repressor that downregulates expression of the *prp* operon and further limiting propionate catabolism [57]. The loss of *ΔprpE* and *ΔprpC* did not have any impact on carbonatogenesis in our screen (data not shown), but while these mutations would prevent the entry of propionate into the methylcitrate cycle, propionate catabolism could proceed via the propionyl-CoA degradation pathway, allowing the release of the Ca^2+^ ion for carbonatogenesis [58]. An acetyl-CoA ligase (*acs*) can also compensate for the loss of the propionyl-CoA *prpE* [59]. Although there may be additional complexity in the propionate catabolism pathway that was not captured in this assay, the data support the model that catabolism of the calcium carboxylate is critical.

Our prior work in *Salmonella* also demonstrated an important role for calcium transport in carbonatogenesis, with a knockout in the Ca^2+^ transporter ChaA having a dramatic effect on precipitation; however, we did not identify *ΔchaA* during the Keio library screen [27]. Research has shown that in *E. coli* ChaA has a higher affinity for K^+^ than Ca^2+^ ions and plays an important role in cell growth under alkaline conditions [60, 61]. We did attempt to use *chaA* as a control in our expression assays, but of the few pPRO::*chaA* transformants obtained, all contained amino acid mutations on the cytoplasmic face of the protein compared to the WT *chaA* sequence (data not shown). As pPRO24 is a multicopy plasmid, these data suggested that functional *chaA* is toxic to the cell at levels beyond WT *E. coli,* even on near-neutral (pH 7.2) media [31, 60]. What we did identify was the transporter, *zntB* (Table 1), which has previously been associated with Zn^2+^ transport. While ZntB also binds Cd^2+^ and Ni^2+^, it does not bind Mn^2+^, Mg^2+^ or Cu^2+^, no studies have evaluated Ca^2+^ binding [47, 48]. Overexpression of *zntB* showed a significant increase in carbonate precipitation in both the uninduced and induced treatments compared to the overexpression of CA (Fig 5). No significant difference was detected between induced and uninduced pPRO::*zntB*; however, the P*_prpB_* promoter is leaky and as a membrane protein, ZntB would have a longer half-life than cytoplasmic CA [62]. This could allow ZntB to accumulate in the membrane prior to induction, reducing the observed impact upon induction (Fig 5). Alternatively, we may have exceeded the solute availability for Ca^2+^ or CO_2_ in the reaction, limiting maximal available CaCO_3_ production.

The mutations in central metabolism appeared to affect carbonatogenesis through a combination of growth defects and electron transport (Table 1, Fig 1). The loss of Complex I (*nuoABCEFGHIJK*) dramatically reduced CaCO_3_ precipitation across all substrates tested, even while the pH of the media was similar to WT (Fig 2). Complex I functions to accept electrons from NADH_2_ to reduce ubiquinone in the membrane, pumping four protons out of the cell [63, 64]. The loss of Complex I does not prevent central metabolism from proceeding, as this loss can be compensated by a second NADH oxidoreductase complex (encoded by *ndh*), but this complex lacks the transmembrane component and does not translocate protons across the membrane [65, 66], which could explain why *ΔnuoB* did not dramatically lower extracellular pH (Fig 2A) [67, 68]. Of note is that the onset of carbonatogenesis was delayed even when the pH exceeded the SI for CaCO_3_ (Fig 2C), suggesting that the loss of Complex I may interfere with the onset of CaCO_3_ precipitation in *ΔnuoB*, even when the required physiochemical conditions are met (Fig 2B and C).

We identified another key player in central metabolism that prevented precipitation: SQR (Table 1). A knockout in *ΔsdhC* dropped the pH considerably on all the media examined (Fig 2A). Given that mutations in the *sdh* operon also interfere with electron transport, the reduced pH could be due to increased production of acetate to reduce pH [67, 68]. Given its prominent role in cellular metabolism, the *sdh* operon is regulated by several factors, including the ferric uptake regulator (Fur) protein and we decided to explore the impact of Fe(III) amendment on carbonatogenesis. The Fe(III) amendment accelerated CaCO_3_ precipitation (Fig 4), which could occur through the inactivation of Fur and upregulation of *sdh* expression. Nonetheless, it is unlikely that the B-4mSu media is Fe-limited and the identification of multiple genes involved in iron transport (Table 1) suggests that such impacts may extend beyond the expression of SQR.

In addition to genes involved in electron transport in central metabolism, mutations were also identified in *lhgD* (*ygaF*) and *ubi,* which are involved in electron transport (Fig 1, Table 1). It is possible that limiting the entry of reducing equivalents to the quinone pool promotes overflow metabolism, which would otherwise lead to the production of acids, reducing cellular pH and MICP [67, 68]. Alternatively, the enzyme LhgD converts 2-hydroxyglutarate to α-ketoglutarate in the catabolism of lysine (Fig 1), and the loss of this gene could affect amino acid metabolism and the release of NH_4_^+^ into the media. Given the need for amino acids in the precipitation media, it suggests that this NH_4_ ^+^ release plays an important role in raising pH and promoting carbonatogenesis.

Collectively, these results allow us to refine our MICP model (Fig 6), which may be more dependent on CO_2_ in the cellular environment than previously recognized [27]. Our original model requires the cell to take up Ca^2+^ from the catabolism of a calcium-rich carbon source, which is confirmed by the identification of *prpD* in our assay (Table 1; Fig 2). The uptake of excess Ca^2+^ requires its transport outside of the cell and in *Salmonella* this role was satisfied by ChaA, but in *E. coli* it appears to be driven by ZntB (Fig 5). We had originally believed that the immobilization of the excess Ca^2+^ outside of the cell as CaCO_3_ was an active process, with CA generating the CO_3_ ^2-^ ion for precipitation from atmospheric CO_2_ [27]. While the role of CAs in this process was confirmed in this work (Fig 5), we believe this is actually a passive process based on the metabolic need for CO_2_ that consequently alters the conditions surrounding the cell to promote MICP.

In the MICP process using Ca^2+^ homeostasis occurs with all calcium sources tested, but is most pronounced with calcium acetate (in non-*E. coli* species) and calcium succinate [26, 27, 30]. Both acetate and succinate catabolism bypass the CO_2_ liberated by the decarboxylation of α-ketoglutarate and succinyl-CoA (Fig 6). In the case of acetate, the two-carbon acetate prevents regeneration of oxaloacetate, and the cell uses the glyoxylate shunt to lyse isocitrate into succinate, requiring the acetate to be added to the glyoxylate via a condensation reaction to produce malate (Fig 6). Growth on calcium succinate in the absence of glucose similarly prevents a full TCA cycle. Much of the CO_2_ usually generated via the TCA cycle diffuses out of the cell, but a certain portion is used in anabolic pathways, such as lipid synthesis. Under such conditions, growth on calcium acetate is dependent on functioning CA to compensate for the lack of metabolic CO_2_ produced [27]. A significant increase in MICP is also seen when *E. coli* expresses the CA genes *can* and *cynT* (Fig 5) on calcium succinate.

Finally, it has been well established that cellular polymeric structures play a role in MICP, and precipitated CaCO_3_ morphology has previously been shown to be directly affected by cell surface properties [69, 70]. This precipitation appears to be driven by the initiation of nucleation of CaCO_3_ crystals on the cell surface, as mutations impairing this initial phase result in reduced MICP [71]. Such studies have shown that repeating polymers, such as peptidoglycan, teichoic acids, lipopolysaccharides (LPS), and exopolysaccharides (EPS) are able to coordinate bound Ca^2+^, enabling CaCO_3_ crystallization to proceed at lower chemical pressures and promoting precipitation [70–73]. Our work similarly suggests that such structures play an important role in *E. coli,* including peptidoglycan, LPS, and EPS, but expands these polymers to include fimbriae and flagella (Table 1).

The conditions that promote MICP in *E. coli* (CO_2_ uptake, Ca^2+^ enrichment, increased pH, nucleation) actually reflects the conditions in caves that lead to secondary precipitation of CaCO_3_ (such as stalactites and stalagmites) [74]. The bacterial species used to identify this novel MICP pathway were also originally cultured from caves [27]. Secondary deposits form in caves when surface water becomes saturated with CO_2_ as it passes through the soil, creating H_2_CO_3_. As this weak acid dissolves limestone rock (CaCO_3_), it becomes saturated with Ca^2+^ and CO_3_ ^2-^ as it moves through the vadose zone [74]. As this water drips into the atmosphere of the cave, the CO_2_ off-gasses, reducing the H_2_CO_3_. This raises the pH and dramatically increases the SI for Ca^2+^ and CO_3_^2-^, leading to precipitation on nucleation sites provided by existing CaCO_3_ minerals [74]. Our bacterial system is the spatial inverse of this precipitation (Fig 6): the CO_2_ is taken up by the cell via CA to compensate for the reduced cytoplasmic CO_2_, removing H_2_CO_3_ in the microenvironment around the cell. The Ca^2+^ levels also rise through export from the cell. As this process is increasing the availability of CO_3_ ^2-^, amino acid metabolism also increases the pH through the production of NH_4_ ^+^. The sum of these processes leads to a dramatic increase in the SI for CaCO_3_ in the microenvironment surrounding the cell, where polymeric structures promote nucleation and precipitation (Fig 6).

These improvements in our existing model underscore the intricate relationship between microbial metabolism, structure, and mineral nucleation [27]. It also identifies a myriad of targets in *E. coli* that could be altered to increase the CaCO_3_ yield, including: increasing the cellular uptake of CO_2_; increasing the export of Ca^2+^; increasing the generation of NH_4_ ^+^ through amino acid metabolism; and enriching the presence of MICP-promoting polymers around the cell. The consumption of CO_2_ in this process, either by incorporation into the growing CaCO_3_ crystals or in cell biomass, reduces dissolved CO_2_ in the media, which is replaced by equalization with atmospheric CO_2_ [27]. This suggests that if a Ca^2+^ homeostasis system could be engineered (through genetic modification and established conditions) where microbial/mineral CO_2_ uptake exceeds the atmospheric diffusion rate, the reactions could be supplemented with additional CO_2_ to provide a novel carbon capture approach. Nonetheless, the addition of CO_2_ would need to be facilitated in such a way that H_2_CO_3_ does not reduce the pH and knockout MICP, increasing the potential for this method to be utilized in the production of industrially relevant carbonates without the need for ureolysis.

## Acknowledgements

This work was supported by the Defense Advanced Research Projects Agency: Engineered Living Materials grant HR0011-18-9-0007 and NSF grant IIP-2122799.

